# Enhancers associated with unstable RNAs are rare in plants

**DOI:** 10.1101/2023.09.25.559415

**Authors:** Bayley R. Mcdonald, Colette Picard, Ian M. Brabb, Marina I. Savenkova, Robert J. Schmitz, Steven E. Jacobsen, Sascha H. Duttke

**Affiliations:** School of Molecular Biosciences, College of Veterinary Medicine, Washington State University, Pullman, WA 99164, USA; Department of Molecular Cell and Developmental Biology, University of California at Los Angeles, Los Angeles, CA, USA; Department of Genetics, University of Georgia, Athens, GA, USA; Howard Hughes Medical Institute, University of California at Los Angeles, Los Angeles, CA, USA

## Abstract

Unstable transcripts have emerged as markers of active enhancers in vertebrates and shown to be involved in many cellular processes and medical disorders. However, their prevalence and role in plants is largely unexplored. Here, we comprehensively captured all actively initiating (“nascent”) transcripts across diverse crops and other plants using capped small (cs)RNA-seq. We discovered that unstable transcripts are rare, unlike in vertebrates, and often originate from promoters. Additionally, many “distal” elements in plants initiate tissue-specific stable transcripts and are likely *bone fide* promoters of yet-unannotated genes or non-coding RNAs, cautioning against using genome annotations to infer “enhancers” or transcript stability. To investigate enhancer function, we integrated STARR-seq data. We found that annotated promoters, and other regions that initiate stable transcripts rather than unstable transcripts, function as stronger enhancers in plants. Our findings underscore the blurred line between promoters and enhancers and suggest that cis-regulatory elements encompass diverse structures and mechanisms in eukaryotes.

## Introduction

The discovery of rapidly degraded and often unprocessed RNAs, such as enhancer-associated RNAs in mammals (De Santa et al. 2010; Kim et al. 2010), has sparked the ongoing endeavor to demystify their role and potential functions. Methods that capture actively transcribed or “nascent” RNA rather than steady-state transcript levels that are a result of many processes, including initiation, elongation, maturation, and decay (Wissink et al. 2019; Yamada and Akimitsu 2019) were instrumental to this research. These approaches have revealed that unstable RNAs are highly prevalent in vertebrates and are involved in many cellular processes and medical disorders (Palazzo and Lee 2015). Unstable transcripts have also been shown to impact gene expression by interacting with transcription factors (TFs), co-factors, or chromatin (Flynn et al. 2011; Lam et al. 2013; Mousavi et al. 2013; Schaukowitch et al. 2014; Benner et al. 2015; Oksuz et al. 2023), and influence the three-dimensional structure of the genome (Lewis et al. 2019).

Distal, bidirectional unstable transcripts, often referred to as enhancer RNAs (eRNAs), have emerged as preferred markers of active regulatory regions in vertebrates (Kim et al. 2010; Azofeifa et al. 2018; Ding et al. 2018; Arnold et al. 2019). These eRNAs are commonly short, non-polyadenylated, unstable, and generated from bidirectionally transcribed loci (Arnold et al. 2019) although some eRNAs were shown to be spliced or polyadenylated eRNAs (Ørom et al. 2010; Gil and Ulitsky 2018; Arnold et al. 2019). Plants no doubt leverage distal cis-regulatory regions, including traditional enhancers (Timko et al. 1985; Oka et al. 2017; Lu et al. 2019; Ricci et al. 2019) but the prevalence and potential roles of unstable transcripts is largely unexplored (Hetzel et al. 2016; Weber et al. 2016; Yan et al. 2019).

Given the importance of plants as the world’s primary food source and their central role in enlivening and sustaining the environment, it is critical to address this gap in our knowledge. However, high quality nascent RNA sequencing datasets, and especially nascent transcription start site (TSS) data, from plants are currently rare. Although some groups, including ours, have demonstrated nascent RNA sequencing methods are feasible in plants, including GRO-seq (Hetzel et al. 2016; Liu et al. 2018; Zhu et al. 2018), PRO-seq (Lozano et al. 2021) and pNET-seq (Zhu et al. 2018; Kindgren et al. 2020), their application is challenging. Plant cell walls, abundant plastids, and secondary metabolites hinder the necessary isolation of pure nuclei and complicate immunoprecipitation steps. Additionally, plants have five or more eukaryotic RNA polymerases, as well as multiple phage-like and plastid-encoded prokaryotic RNA polymerases (Zhou and Law 2015), and traditional GRO-seq (Core et al. 2008) and PRO-seq (Kwak et al. 2013) methods capture nascent transcripts from all of these RNA polymerases nonspecifically, complicating data interpretation (Liu et al. 2018). Thus, nascent RNA sequencing methods drastically advanced our understanding of unstable transcripts in animals and fungi (Core et al. 2008; Core et al. 2014; Mikhaylichenko et al. 2018; Core and Adelman 2019; Wissink et al. 2019), but less in plants.

Some of the technical limitations described above can be alleviated by exploiting the 5’ cap and specifically enriching for nascent RNA polymerase II transcripts and their transcription start sites (TSSs) (5’GRO, (Lam et al. 2013), also known as GRO-cap (Core et al. 2014). Selective sequencing of 5’ capped ends also increases the sensitivity of these methods to detect short, rare, and unstable transcripts (Duttke et al. 2019; Yao et al. 2022) such as eRNAs (De Santa et al. 2010; Kim et al. 2010; Lam et al. 2013) and promoter-divergent unstable transcripts (Seila et al. 2008). We recently developed capped small (cs)RNA-seq, which leverages these advances and directly enriches for nascent RNA polymerase II TSSs without the need for nuclei isolation, run-on, or immunoprecipitation (Fig. 1a, (Duttke et al. 2019). csRNA-seq is a simple, scalable and cost-efficient protocol that uses 1-3 ug of total RNA, rather than purified nuclei, as input, and is compatible with any fresh, frozen, fixed, or pathogenic species or tissues (Lim et al. 2021; Branche E 2022; Duttke et al. 2022b; Lam et al. 2023). Recently, csRNA-seq was shown to effectively detect eRNAs in human lymphoblast cells (Yao et al. 2022).

**Fig. 1.**
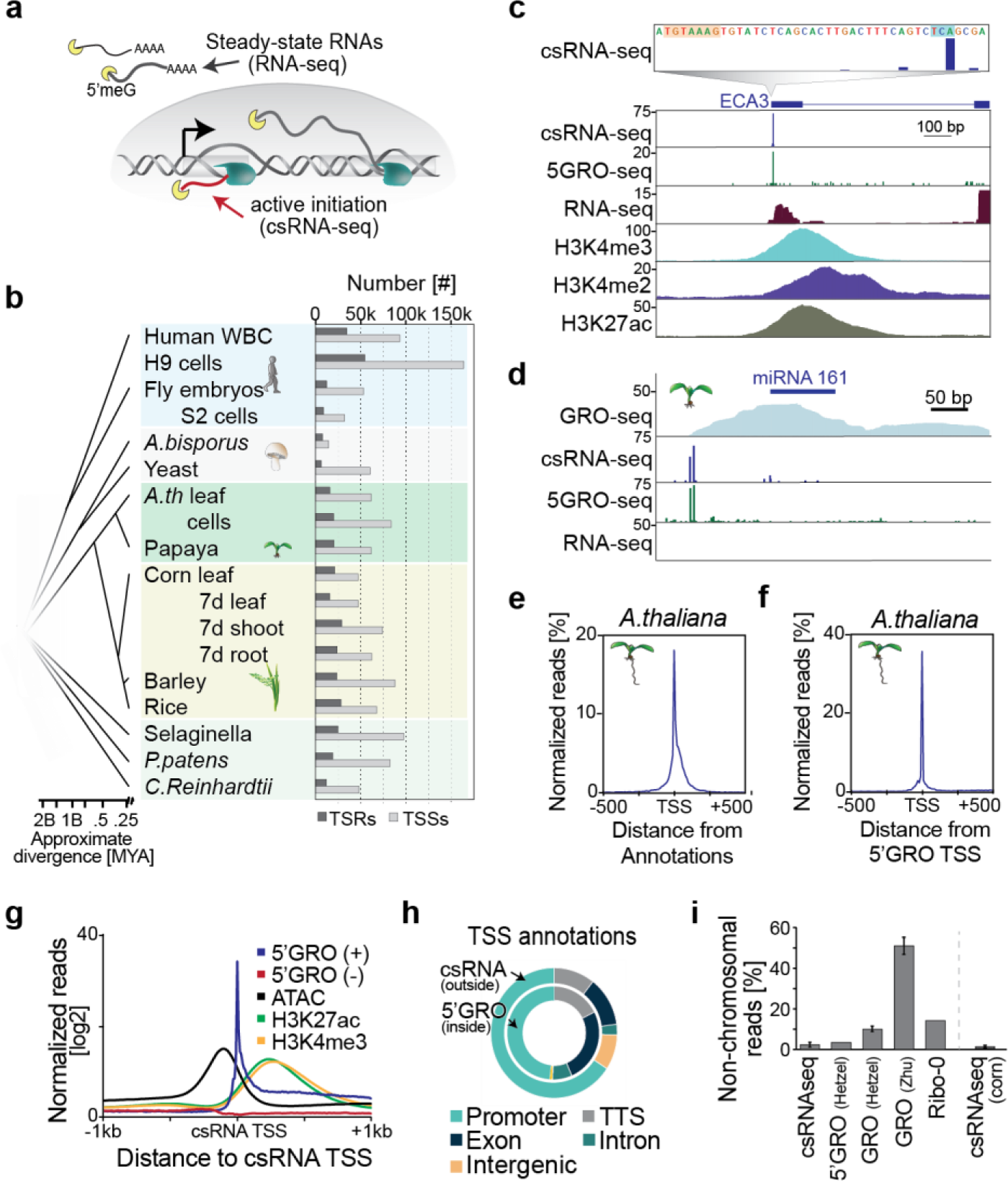
A comprehensive atlas of nascent plant transcription initiation. **a**, Schematic of steady state RNA, as captured by RNA-seq, and actively initiating or nascent transcripts, captured by csRNA-seq. **b**, Overview of samples studied with the numbers of captured transcription start regions (TSRs), which include promoters and enhancers, and of transcription start sites (TSSs). Samples generated in this study are marked with an asterix (*). WBC = white blood cells. **c**, *A. thaliana* ECA3 loci with csRNA-seq at single nucleotide resolution and zoomed out. 5’ GRO-seq and histone ChIP-seq data. **d**, *A. thaliana* miRNA 161 cluster. **e**, Distribution of *A. thaliana* csRNA-seq data from leaves relative to TAIR10 TSS annotations. **f**, Distribution of csRNA-seq TSSs from *A. thaliana* leaves relative to 5’GRO-seq TSSs mapped in 7d-old seedlings. **g**, Distribution of 5’ GRO-seq reads as well as open chromatin (ATAC-seq) and histone marks H3K4me3 and H3K27ac relative to csRNA-seq TSSs in *A. thaliana.* **h**, Comparison of annotations of TSSs mapped by 5’GRO-seq and csRNA-seq in *A. thaliana.* **i**, Percent of non-chromosomal RNA reads captured by csRNA-seq, GRO-seq (Hetzel et al. 2016; Zhu et al. 2018)et, 5’GRO-seq (Hetzel et al. 2016), or total RNA-seq (Ribo0) in *A. thaliana* and maize (csRNA-seq only). These RNAs are not synthesized by RNA polymerase II or other eukaryotic RNA polymerases.

Here, we used csRNA-seq to decipher the prevalence, location, and traits of stable and unstable transcripts across different plant tissues, cells, and species. Our data suggest that vertebrate-like eRNAs are effectively absent in plants. Instead, promoters were the major source of unstable transcripts and intriguingly, promoters and open chromatin regions rather than sites initiating unstable transcription, showed the strongest enhancer activity in the STARR-seq assay.

## Results

### A comprehensive atlas of nascent transcripts in plants

To comprehensively capture nascent transcripts in plants, we performed csRNA-seq on 12 samples from eight plant species chosen for their agricultural and scientific importance (Fig. 1b, Table S1). For comparison, we also performed csRNA-seq on S2 cells from fruit fly (*Drosophila melanogaster*) and integrated published data from fly embryos (Delos Santos et al. 2022), rice (*Oryza sativa*, adult leaves) (Duttke et al. 2019), human white blood cells (WBCs) (Lam et al. 2023), human lymphocytes (Duttke et al. 2019), and fungi (common mushroom, *Agaricus bisporus*; yeast, *Saccharomyces cerevisiae*) (Duttke et al. 2022a) (Fig. 1b).

csRNA-seq accurately captured actively transcribed stable and unstable RNAs and their transcription start sites (TSSs) genome-wide and at single nucleotide resolution, as exemplified by the *A. thaliana* ER-type Ca2+ ATPase 3 (ECA3) locus (Fig. 1c) or unstable pri-miRNA 161 (Fig. 1d), similar to other nascent methods but, on average, with less background noise (Fig. 1c,d, Fig. S1a). As expected, csRNA-seq-captured TSSs were enriched in proximity to annotated TSSs genome wide (Fig. 1e, Fig. S1b-e). About one third of the csRNA-seq TSSs mapped in *A. thaliana* leaves were identical to those mapped by 5’ GRO-seq in 7-day old seedlings, and nearly all were within 200 bp (Fig. 1f (Hetzel et al. 2016)). TSSs identified by csRNA-seq were also similar to those we identified by 5’ GRO-seq in *Physcomitrium patens, Chlamydomonas reinhardtii* and *Selaginella moellendorffii* (Fig. S1f-h). Thus, csRNA-seq accurately captures actively initiated transcripts and their TSSs in diverse plant species and tissues.

To validate our csRNA-seq TSSs further, we examined their association with the chromatin and epigenomic landscape (Ricci et al. 2019; Zhao et al. 2022; Wang et al. 2023). As expected for active TSSs, chromatin accessibility (ATAC-seq) peaked just upstream of csRNA-seq-captured TSSs in both *A. thaliana* (Fig. 1g) and maize (Fig. S1i). Histone modifications associated with transcription initiation, such as H3K27ac and H3K4me3 (Creyghton et al. 2010; Lauberth et al. 2013), were found downstream of csRNA-seq TSSs (Fig. 1g, Fig. S1i). Regions of transcription initiation were also enriched in genomic regions annotated as associated with transcription and were mainly found at promoter regions (Fig. 1h). Sites of transcription initiation across plant species revealed a similar pattern to *A. thaliana*, with the majority of TSSs located within annotated promoter regions (Fig. S2, Table S2). Additionally, csRNA-seq exhibited efficient and specific enrichment of 5’ capped RNA polymerase II transcripts, with only a small percentage of reads mapping to non-chromosomal regions like plastids or mitochondria (Fig. 1i). These data further support that csRNA-seq accurately identifies nascent transcripts and their TSSs from total RNA across diverse plant species.

In eukaryotes, most genes display dispersed transcription initiation from multiple TSSs within 20-100 bp in the same promoter or enhancer region (Haberle and Stark 2018; Murray et al. 2022). Therefore, we will hereafter jointly refer to all strand-specific individual or clusters of TSSs as transcription start regions (TSRs, Fig. 2a (Luse et al. 2020)). This approach also avoids implying functionality of identified regions beyond initiating transcription (Halfon 2019), and definition of “enhancers” based on annotations, whose quality varies strongly across the species or even for different cell types within a species (Shamie et al. 2021).

**Fig. 2.**
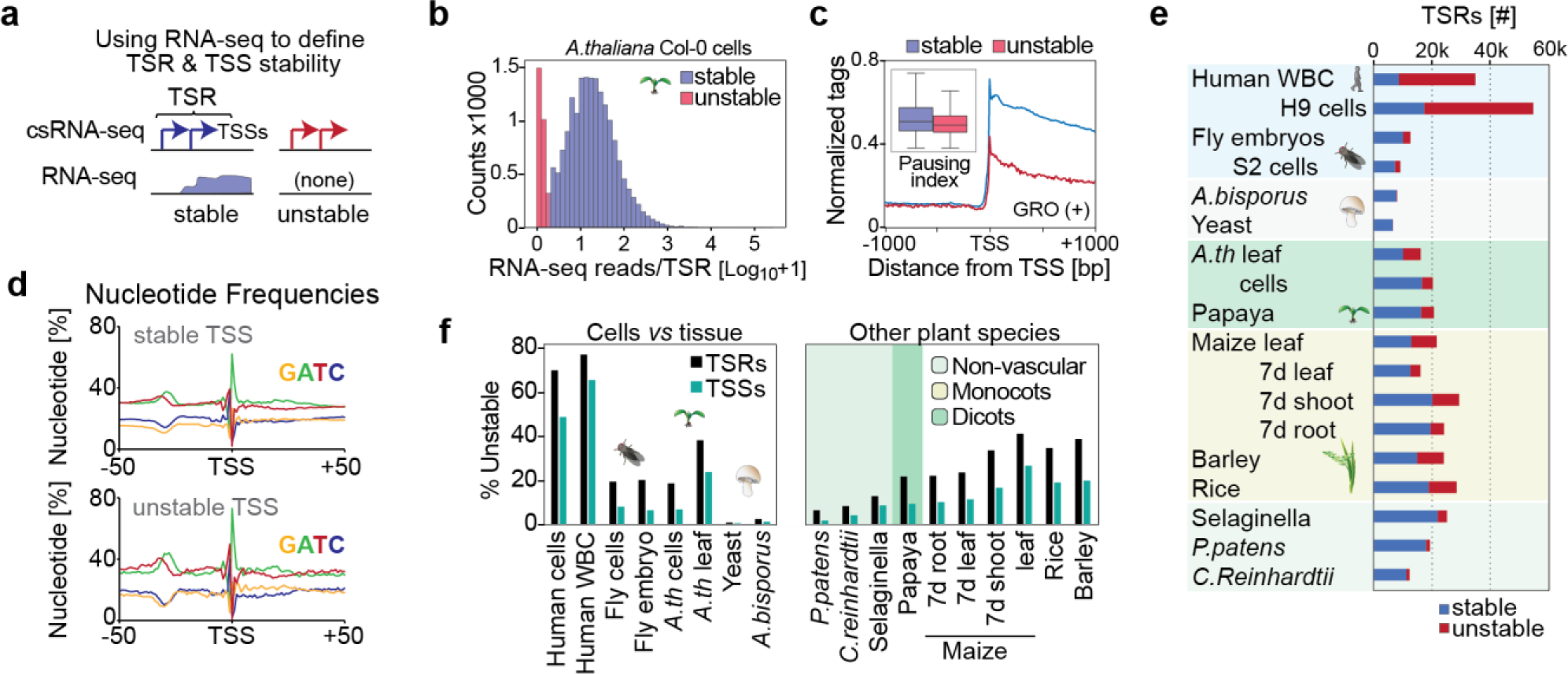
Unstable RNAs are infrequent in plants. **a**, Schema of how transcript stability was determined by integrating total RNA-seq read counts from -100 to +500 with respect to the major TSSs within TSRs identified by csRNA-seq. **b**, Distribution of RNA-seq reads per million within -100 to +500 bp relative to the main TSS of each TSRs, plotted as [log10 +1]. **c**, GRO-seq signal (+ strand only) in *A. thaliana* (Hetzel et al. 2016; Zhu et al. 2018) in proximity to the TSS of stable and unstable transcripts. Inset: calculated pausing index (reads -100, +300 / reads +301, +3000, see methods). **d**, Metaplot of nucleotide frequency with respect to the +1 TSS as defined by csRNA-seq for stable and unstable transcripts in *A. thaliana* **e**, Summary of the number of stable and unstable TSRs in each sample analyzed. **f**, Percent of TSRs and TSSs initiating unstable transcripts across all species and tissues assayed.

The number of detected TSRs and TSSs varied from about 6.5k TSRs with 60k TSSs in yeast, to about 60k TSRs and 165k TSSs in human H9 lymphocytes (Fig. 1b, Table S1). Among plant species, we observed a range of TSRs and TSSs, from 12.6k TSRs with 48k TSSs in *C. reinhardtii* to 30k TSRs and up to 88k TSSs in some monocots (e.g. barley). In total, we identified >380,000 TSRs with over 1.25 million TSSs. This comprehensive atlas provides a valuable resource for studying transcription and gene regulation in plants and spans over 1.5 billion years of evolution.

### Unstable RNAs are infrequent in plants

CsRNA-seq captures active transcription initiation and thus both stable and unstable RNAs. To infer transcript stability, we took advantage of total RNA-seq, which reports stable, steady-state RNAs. Specifically, we estimated transcript stability by quantifying total RNA-seq reads near csRNA-seq TSSs (Fig.2 a)(Blumberg et al. 2021). This approach is independent of genome annotations, which vary drastically in quality among the studied samples. TSSs of unstable RNAs have few-to-no strand-specific RNA-seq reads downstream (e.g., Fig. 1d), whereas stable RNAs are readily detected by RNA-seq (e.g., Fig. 1c, (Duttke et al. 2019). The number and percentage of TSRs and TSSs that give rise to unstable transcripts (uTSRs and uTSSs, respectively) is dependent on peak csRNA-seq calling cutoff and total RNA-seq coverage (Fig. S3a). We required unstable RNAs to have less than 2 per 10 million RNA-seq reads within -100 bp to +500 bp of the major TSSs within the TSR, in line with the observed bimodal distribution of TSR stability (Fig.2 b).

The number of stable TSRs was with ∼7-21k comparatively similar across kingdoms. However, the number and percentage of uTSRs and uTSSs varied up to 100-fold (Fig. 2e, Table S1). In humans, up to 75% of all TSRs were uTSRs, whereas in fruit flies, this frequency was about 20%, and in the fungi yeast and *A. bisporus*, it was less than 2% (Fig. 2e, Table S1). In plants, the percentage of uTSRs ranged from 6% to 40%, with uTSRs being more common in monocots (e.g. rice, maize, barley) compared to dicots (e.g. *A. thaliana*, papaya) or nonvascular plants (e.g. Selaginella, *P. patens*). There was also variability in the proportion of uTSRs among different tissues within the same organism, for example in different maize tissues (Fig.2e, Fig. S3c). Importantly, these numbers likely present the upper limit of unstable transcripts. CsRNA-seq is orders of magnitude more sensitive than RNA-seq (Duttke et al. 2019); as a result, recently activated or tissue-specific TSRs might be readily detected by csRNA-seq, but not total RNA-seq. Thus, these TSRs that are in fact stable could be misclassified as unstable RNAs. To minimize resulting bias, we therefore focused our analysis where possible on tissues in near-quiescent states, like mature leaves and cultured cells. However, it is likely that the true number of uTSRs is even lower than what we are reporting.

Unstable transcripts could also result from premature termination prior to RNA polymerase II pause release (Core and Adelman 2019). Because csRNA-seq alone cannot discern between this scenario and rapid degradation post RNA processing (canonical “instability”), we integrated published GRO-seq data from *A. thaliana* leaves and seedlings (Zhu et al. 2018; Hetzel et al. 2016) and estimated pausing. Plotting read density within 1kb of TSSs and calculating the pausing index (reads -100, +300 / reads +301, +3000, relative to TSSs (Chen et al. 2015)) showed only a modest increase in proximal RNA polymerase II occupancy near the TSSs of unstable transcripts compared to stable ones (Fig. 2c). Thus, consistent with the absence of canonical promoter-proximal pausing in plants (L. Core and Adelman 2019), pausing-dependent termination is unlikely to explain most of the unstable transcripts identified in this study.

Importantly, although unstable transcripts were on average more weakly initiated than stable ones (Fig. S3b), the DNA sequence composition surrounding TSRs initiating stable and unstable transcription was highly similar (Fig. 2d). TSRs of both groups had hallmarks of canonical cis-regulatory elements including a TATA-box and Initiator core promoter signature, emphasizing that these unstable TSRs are not just transcriptional noise. Furthermore, *de novo* motif analysis of sequence motifs in proximity to TSSs (-150, +50, relative to the TSS) initiating stable or unstable transcripts also revealed highly similar occurrences of transcription factor binding sites (r>0.95) for all samples with notable uTSS (>1.5k, Fig. S4). These results not only emphasize that both stable and unstable TSSs captured by our method are *bona fide* TSSs, but also suggest that similar regulatory mechanisms support the initiation of stable and unstable transcripts in plants.

Unstable transcripts often display cell-type-specific expression (Arnold et al. 2019), which may compromised their detection in complex samples. To address this notion, we compared the detection of uTSRs from clonal cell populations versus tissues. About 18% of all detected TSRs were uTSRs in *A. thaliana* Col-0 cells, compared to 37% in leaves. By contrast, 19% and 20% of TSRs were unstable in fruit fly S2 cells and in 0-12h embryos, respectively; 0.5% versus 2% were unstable in single cell yeast versus the multicellular mushroom *A. bisporus*, and 68% and 75% were unstable in human lymphocytes versus white blood cells (Fig. 2f). Thus, a higher percentage of uTSRs and uTSSs was observed in complex tissues across all kingdoms (Fig. 2f, Fig. S3c, Table S1). Together, these data argue that the previously reported underrepresentation of unstable TSRs and RNAs in plants (Hetzel et al. 2016) is not due to their limited detectability in complex tissues. Indeed, our data consistently captured uTSSs and uTSRs in diverse plant species, fruit flies and fungi, but they are much less prevalent in all these organisms than in humans.

### Origins of plant unstable transcripts

Studies in vertebrates have identified many distinct classes of unstable RNAs, such as short, bidirectional eRNAs, promoter-divergent transcripts, and others (Seila et al. 2008; Almada et al. 2013; Kim and Shiekhattar 2015; Field and Adelman 2020). As genomic locations of origin, rather than functional assays, were often used to classify these transcript types, we compared the genomic locations of uTSRs in *A. thaliana* Col-0 cells and human lymphocytes, where quality annotations are available. In total, we found 3,651 uTSRs in *A. thaliana*, compared to 37,315 in humans. While this number is about the same when normalizing for genome size, it is important to remember that with 16,527 vs. 17,268, a similar number of stable transcripts was expressed in both species.

While unstable transcripts from promoter divergent or antisense transcription were prominent in humans, unstable transcripts in plants predominantly originated from promoters in sense (Fig. 3a, b). About 27% of uTSRs in *A. thaliana* were initiated in the sense orientation from annotated promoters, compared to 17.8% in humans (Fig. 3a). These promoters in *A. thaliana* were often tissue-specific and did not share specific pathways or gene sets (Fig. S3e), hinting that the observed initiation of uTSRs may be sporadic, rather than resulting from a common or regulated mechanism. Approximately 7.3% of uTSRs were promoter-proximal and divergent, compared to 15.3% in human lymphocytes (Fig. 3a, Fig. S3d). Another 1.5% and 5.4% in *A. thaliana*, and humans respectively, were within 300 bp downstream of the TSS and therefore TSS antisense. We found that 2.7% of human uTSRs and 5.8% in *A. thaliana* annotated to single exon transcripts like snRNA and snoRNA. These short transcripts are inefficiently captured by total RNA-seq due to their small size, and therefore may not, in fact, be unstable (Duttke et al. 2019). Some uTSRs were found in the proximity of genes encoding miRNAs (Fig. 3a), thus likely pri-miRNA promoters, and only 2.7% of human uTSRs and 5.8% *A. thaliana* uTSRs were in exons.

**Fig. 3.**
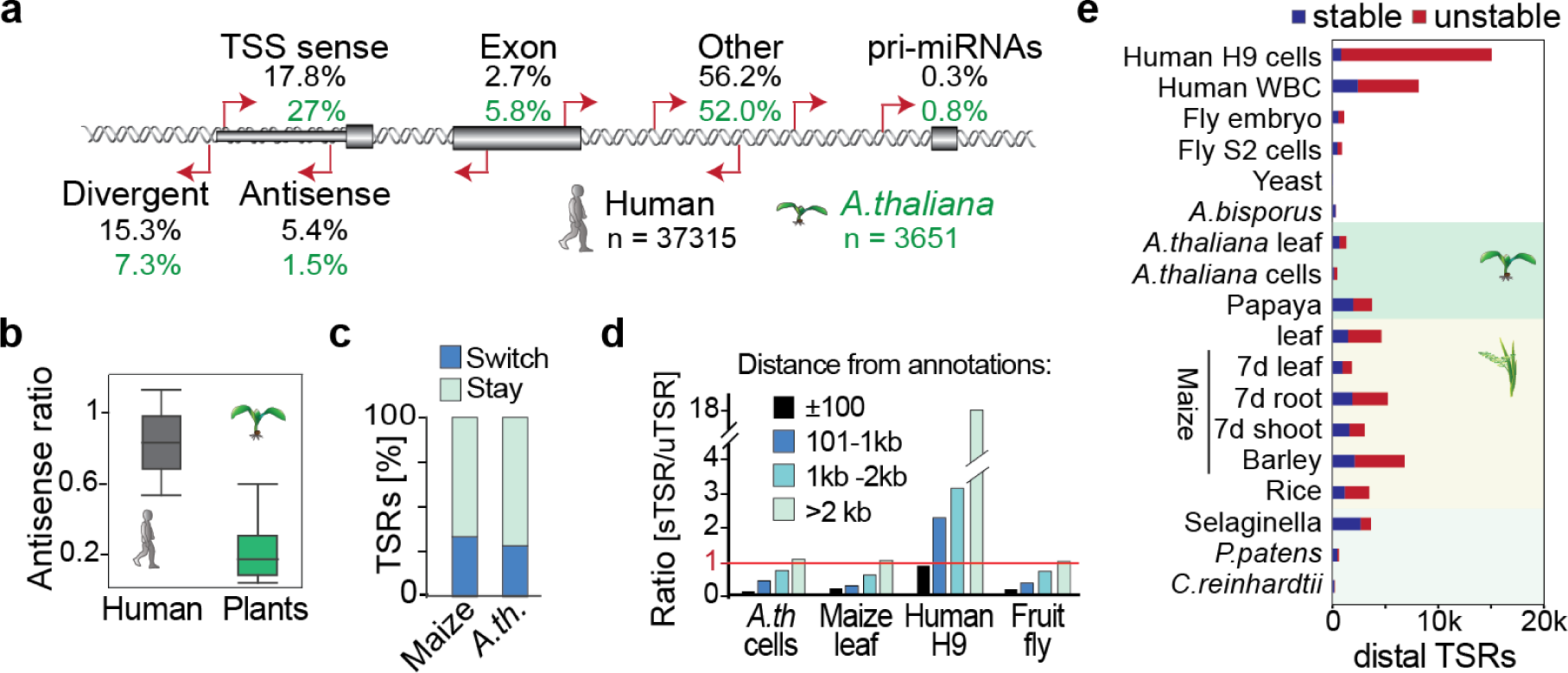
Distinct origins of stable and unstable transcripts in humans, plants, and other species. **a**, Classification of uTSR genomic sites in human H9 lymphoblast and *A. thaliana* Col-0 cells, relative to current annotations (Araport11/ gencode.42). TSS = ±275 bp of 5’ gene annotation in sense direction; TSS antisense, within the TSS region but antisense; TSS divergent, initiating from -1 to -275bp to the TSS. **b**, Ratio of promoter-proximal antisense transcription reveals most plant but not human unstable transcripts to initiate in *sense* direction. Ratio of TSRs in antisense to genome-annotated TSSs (-275 to +275 relative to the annotated TSS) divided by the number of total TSRs that mapped to annotated TSS. **c**, Percentage of TSRs that switch between initiating stable and unstable transcripts among *A. thaliana* Col-0 cells and leaves, maize adult leaves and 7d-old leaves, shoot, and roots, and Human H9 cells and white blood cells (WBCs). **d**, Number of TSRs initiating unstable divided by stable transcripts relative to distance to genome annotations by regions (*i*) ± 100 bp, (*ii*) 101-1,000 bp, (*iii*) 1,001-2,000 bp, (*iv*) > 2,000 bp for *A. thaliana,* fruit fly S2 and human lymphoblast cells as well as maize leaves. **e**, Number TSRs >2,000 bp from annotations that initiate stable or unstable transcripts.

Therefore, most uTSRs initiated outside of annotated regions in both human lymphocytes (56.2%, ∼21,000 uTSRs) and Col-0 cells (54.4%, ∼1950 uTSRs) (Fig. 3a). It is important to reiterate that, given the higher sensitivity of csRNA-seq over RNA-seq (Duttke et al. 2019), many of these sense transcripts classified as unstable could be newly activated genes or non-coding RNAs, suggesting the true number of unstable RNAs found in plants to be even lower than what we are reporting. However, as detailed below, many of these “distal loci” in plants but not humans also initiated stable transcripts in other tissues.

### Many plant promoter and distal TSRs give rise to stable and unstable transcripts

To determine if TSRs can switch between initiating stable and unstable transcripts, we compared their stability in *A. thaliana* Col-0 cells and leaves, as well as the young and adult leaf, shoot and root samples from maize. We found that about 28.4% of TSRs in *A. thaliana* and 33.4% in maize switched from unstable to stable or vice versa in the different samples (Fig. 3c). Moreover, about 18.5% of transcripts switched between stable and unstable in biological replicates of the same sample. Thus, many TSRs give rise to stable transcripts in one tissue context and unstable transcripts in another.

Given these findings, we also explored the spatial relationship of stable TSRs relative to annotations across species. In *A. thaliana* cells, maize leaves and S2 cells from fruit fly, TSRs within 100 bp of annotations were predominantly stable, but about 28% of all uTSRs in *A. thaliana* cells, 50% in maize leaves, and 64% in S2 cells also located to these promoter regions (Fig. 3d, Fig. S3f). In human, a comparable number of stable and unstable transcripts initiated from within 100 bp of annotated promoters. However, while in most of the samples that we analyzed, a similar or even higher number of TSRs >100 bp of annotated promoters initiated stable transcripts compared to unstable transcripts (Fig. 3d,e), distal TSRs in human samples predominantly initiated unstable transcripts. We note that the total number of distal TSRs initiating stable transcripts was similar in human and plant samples, whereas the number of distal TSRs initiating unstable transcripts was higher in human samples, particularly lymphocytes (Fig. 3e).

Thus, while the distance to annotations and the distal-proximal classification depends on genome size and annotation quality, distal regions were consistently enriched for uTSRs in humans, but not in the other species investigated. In total, we identified 19,397 distal TSRs in plants that initiated stable RNAs. Overall, these findings caution against classifying distal TSRs as uTSRs. Indeed, many distal TSRs initiate stable RNAs in plants, and thus may be promoters of unannotated genes or non-coding RNAs further alluding to unannotated promoters and cell type-dependent stability as the major source of unstable transcripts in plants.

### Canonical vertebrate enhancers are rare in plants

Most human promoters and enhancers start transcription in both forward and reverse directions, often from distinct core promoters (Core et al. 2014; Duttke et al. 2015). By contrast to this predominantly bidirectional nature of transcription initiation in humans, transcription was largely initiated unidirectionally in plants, flies and fungi (Fig.4 a,b, Fig. S5a). On average, only 4.7% of TSRs in plants initiated bidirectional unstable transcripts, most of which were promoter-proximal (Fig.4a,b). For instance, in *A. thaliana* leaves, 62% and 91% of bidirectional TSRs were within 100 bp and 2 kb of annotated 5’ ends respectively (Fig. S5b,c).

**Fig. 4.**
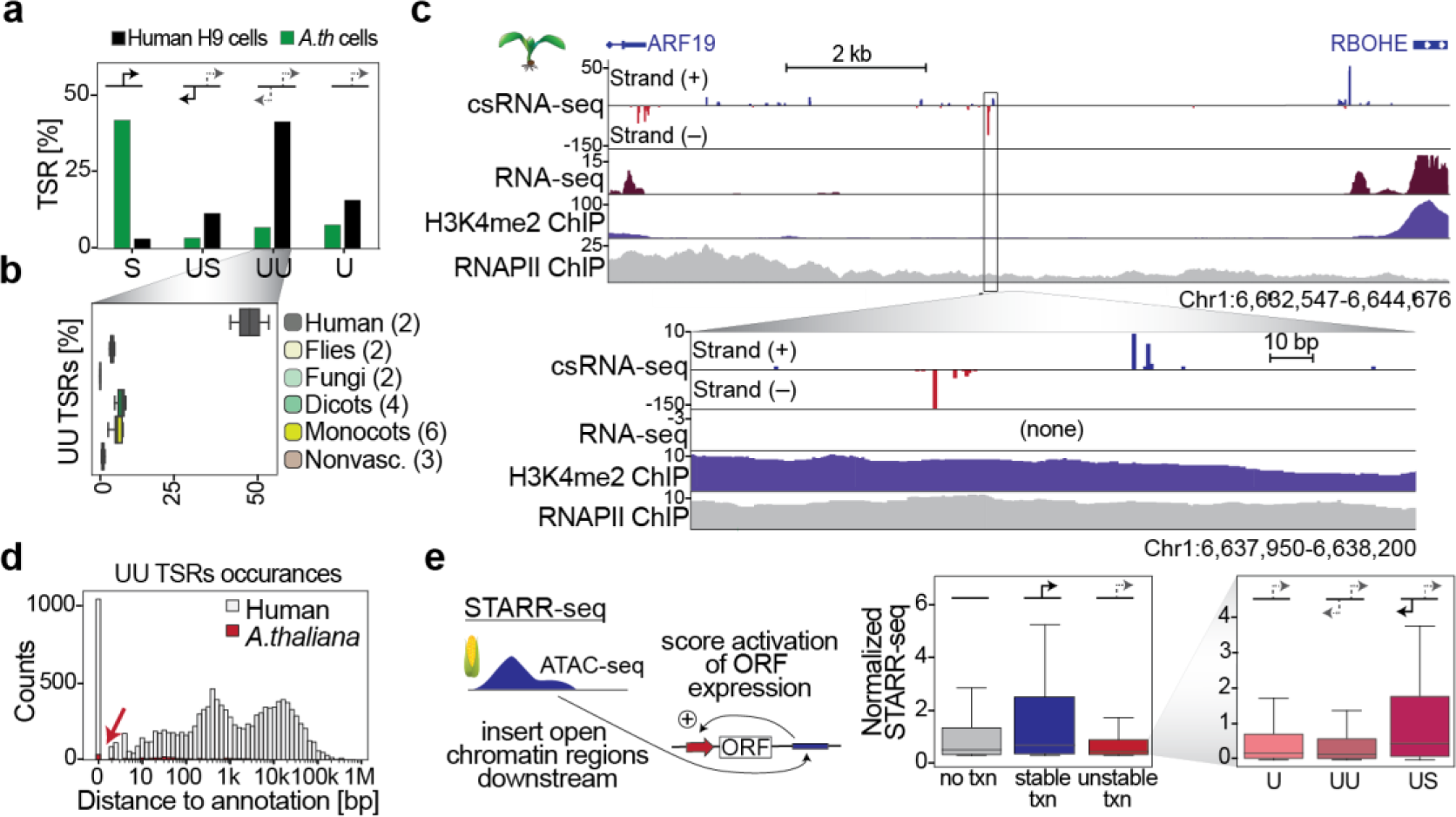
Vertebrate-like enhancers are rare in plants and have less enhancer activity than promoters. a, Overview of TSR directionality and type in human H9 lymphoblast and *A. thaliana* Col-0 cells.; S, TSR is stable and unidirectional; US, TSR produces an unstable sense transcript and a stable antisense transcript; UU, TSR produces unstable sense and antisense transcripts; U, TSR is unstable and unidirectional. **b**, Average percentage of bidirectional unstable transcription in samples from humans (H9 cells, WBC), flies (*D. melanogaster* embryos, S2 cells), fungi (*S. cereviciae, A. bisporus*), dicots (*A. thaliana* cells and leaf, papaya), monocots (maize [4], rice, barley), and nonvascular plants (Selaginella, *P. patens, C. reinhardtii*). **c**, Example of one of the 72 distal TSRs in *A. thaliana* leaves initiating unstable bidirectional transcription. **d**, Distribution of distance to nearest genome annotations for all TSRs initiating unstable bidirectional transcription; annotations in human H9 and *A. thaliana* Col-0 cells. **e**, Overview of the STARR-seq assay (Arnold et al 2013), left) which measures the ability of DNA regions, here all open chromatin regions in maize captured by ATAC-seq (Ricci et al 2019), cloned downstream of a minimal promoter to enhance its transcription. Enhancer function, as measured by STARR-seq promoter activity [scaled by 100], was subgrouped by csRNA-seq in tissue defined TSR type (no, stable or unstable transcription initiation). Regions initiating unstable transcription were further subgrouped by their initiation styles (U, UU, US).

Although there were some instances of distal bidirectional initiation of unstable transcripts in plants, reminiscent of canonical mammalian eRNAs (Fig.4c), they were rare and potentially too few to serve as reliable markers for plant enhancers. For instance, only 361 (1.8%) and 72 (0.5%) TSRs in *A. thaliana* Col-0 cells and leaves, respectively, initiated distal bidirectional unstable transcripts. By contrast, 9,318 (17%) of TSRs in human H9 lymphocytes initiated bidirectional unstable transcripts that were >2kb of annotated 5’ ends (Fig.4d, Fig. S5d). This difference is not simply due to genome size or gene density: even in monocots with large genomes the number of distal, unstable, and bidirectional initiation events varied from only 400-857 events, contributing to a maximum of 3.2% of TSRs (Fig. S5d). As such, distal bidirectional uTSRs are rare in plants.

### Promoters may be stronger enhancers in plants

To explore the functionality of the distal transcription initiation events that we detected in plants, we generated csRNA-seq data matching published STARR-seq data from maize 7d leaves (Ricci et al. 2019). In this assay, open chromatin regions were cloned downstream of a minimal promoter and their ability to enhance transcription quantified (Arnold et al 2013). The majority (92%) of the csRNA-seq TSRs were covered by the STARR-seq library, indicating effective coverage of the maize genome (Fig. S6a). Notably, we found that TSRs initiating stable transcription (i.e., promoters) showed the strongest enhancer activity in plants. Transcription activity, as assayed by csRNA-seq, was overall positively correlated with STARR-seq enhancer activity (r = 0.49, Fig. S6b). Consistent with these findings, regions with high STARR-seq activity were enriched for binding sites for strong activators like GATA or EBF factors while inactive regions had repressors including RPH1, HHO3, and ARID (At1g76110, Fig. S6c). These findings suggest that the competence of a regulatory element to recruit RNA polymerase II contributes to its enhancer activity, as assessed by STARR-seq. However, most promoters and even more uTSRs showed little STARR-seq enhancer activity (Fig. S6d) and substantial STARR-seq enhancer activity was also observed for many open chromatin regions that were transcriptionally inactive (Fig. 4e), as assayed by csRNA-seq.

While vertebrate enhancers are commonly marked by unstable bidirectional transcription (eRNAs), initiation from the upstream STARR-seq promoter in plants was most strongly enhanced by TSRs that initiated stable RNAs (Fig. 4e), whereas uTSRs had weaker enhancer activity. Bidirectional uTSRs that resemble vertebrate enhancers, on average, exhibited the weakest activity (“UU”, Fig.4e, Fig. S6b). By contrast to flies where bidirectional but not unidirectional promoters were reported to often act as potent enhancers (Mikhaylichenko et al. 2018), both uni- and bidirectional promoters exhibited similar STARR-seq activity (Fig. S6d). Notably, uTSRs that initiated stable transcription upstream also show the highest enhancer activity among uTSRs. Together, these findings underscore the blurred line between promoters and “enhancers”, propose enhancers as a heterogeneous group, and highlight distinct features of plant transcription.

## Discussion

By interrogating nascent transcription initiation across a wide range of organisms, we discovered that unstable transcripts are rare in plants, and in fact also in fruit flies, and some fungi, compared to mammals. Distal bidirectional initiation of unstable transcripts, which is a hallmark of vertebrate enhancers and a predominant source of unstable transcripts and eRNAs, was particularly uncommon in plants. Instead, unstable transcription was predominantly unidirectional in plants (Hetzel et al. 2016), originated from promoters, and we identified numerous distal regulatory elements that initiated stable transcripts, making them *bona fide* promoters. These findings suggest that a considerable portion, if not the majority, of unstable RNAs in plants may arise from promoters of either known or unannotated genes or non-coding RNAs (Mendieta et al. 2021) and caution against using genome annotations to infer transcript stability or define “enhancers”.

This study also provides a notable resource to the scientific community. Aside from a comprehensive collection of nascent TSS data paired with total RNA-seq and small RNA-seq (csRNA-seq input) for an array of plant species, tissues, and cells, it demonstrates that csRNA-seq can help to refine genome annotations (Shamie et al. 2021), readily captures the entire active RNA polymerase II transcriptome in plants and across eukaryotes, and serves as a proof-of-concept how csRNA-seq opens up new opportunities to advance our understanding of gene regulation. For instance, csRNA-seq can be readily applied to investigate nascent transcription in a wide range of scientifically or agriculturally important field samples and tissues, allowing for the decoding of gene regulatory networks implicated in biotic or abiotic stress responses.

Our findings also shed light on the discussion surrounding the role and existence of vertebrate-like eRNAs in plants (Weber et al. 2016; Zhang et al. 2022) and further blur the line between the concepts of canonical promoters and enhancers. While distal loci initiating bidirectional unstable transcripts were found in all plant species studied (Fig. 4, Fig. S5d), they were rare, and in some instances, initiated stable transcripts in other tissues or samples from the same plant. Combining csRNA-seq (Duttke et al. 2019) with STARR-seq (Arnold et al. 2013; Ricci et al. 2019), demonstrated that genomic regions initiating stable transcription function as stronger enhancers in plants than those starting unstable transcripts. Intriguingly, among plant TSRs, those resembling mammalian-like enhancers, defined as initiating bidirectional unstable transcription, exhibited the weakest activating properties by STARR-seq. In addition, it is important to note that the number of distal TSRs initiating unstable transcription are also likely too few to make up all plant enhancers. While enhancers defined by eRNAs vastly outnumber genes in humans (Dean et al. 2021), orders of magnitude more stable than unstable transcripts were observed in plants.

It is further notable that many regions that did not initiate transcription in the plant genome, as assayed by csRNA-seq, exhibited STARR-seq enhancer activity, on average more than uTSRs (Fig. 4e). Furthermore, unidirectional plant promoters, on average, displayed similar enhancer activity than bidirectional ones. Contrasting these observations with findings in mammals (Ding et al. 2018; Arnold et al. 2019) or flies, where bidirectional promoters were reported to often act as potent enhancers while unidirectional promoters generally cannot (Mikhaylichenko et al. 2018), suggests that plant promoters may possess distinct attributes. However, it is also possible that gene regulatory elements form a continuum, and that different species or gene regulatory contexts preferentially leverage different parts of it. While “canonical vertebrate enhancers” with eRNAs may be prevalent in some animals, reports of processed eRNAs (Ørom et al. 2010; Gil and Ulitsky 2018; Arnold et al. 2019), enhancers functioning as context-dependent promoters (Kowalczyk et al. 2012), and the important role of enhancers to serve as promoters in the birth of new genes (Ludwig et al. 2000), speak to such a continuum and “enhancers” as heterogeneous group of regulatory elements (Zentner et al. 2011; Link et al. 2018; Halfon 2019; Panigrahi and O’Malley 2021). If true, this continuum hypothesis would propose that there may also be untranscribed regions or unidirectional promoters that function as “enhancers” in other species including humans.

## Methods

### Plant material and growth conditions

*A.thaliana* Col-0 mature leaves were collected from plants grown as described (Wang et al. 2023), while *A.thaliana* Col-0 suspension cells (Concia et al. 2018) were grown in 250-mL baffled flasks containing 50 mL of growth medium (3.2 g/L Gamborg’s B-5 medium, 3 mM MES, 3% [v/v] Suc, 1.1 mg L−1 2,4-dichlorophenoxyacetic acid). The cultures were maintained at 23°C under continuous light on a rotary shaker (160 rpm) and kindly provided as a frozen pellet by Dr. Ashley M. Brooks. Barley (*Hordeum vulgare*) RNA was isolated by Dr. Pete Hedley from embryonic tissue (including mesocotyl and seminal roots; EMB) isolated from grain tissues 4 days past germination (Mascher et al. 2017). *Physcomitrium* (*Physcomitrella*) *patens* (Gransden) was grown on plates with BCDA medium in a growth cabinet at 21°C under 16h light. *Selaginella moellendorffii* was purchased from Plant Delights Nursery (https://www.plantdelights.com/collections/selaginella/products/selaginella-moellendorffii) and grown at the window under normal daylight for 1 week prior to isolating RNA from stems and leaves. *C. reinhardtii*, which was kindly provided by Dr. Will Ansari and Dr. Stephen Mayfield (UC San Diego), was grown to late logarithmic phase in TAP (Tris–acetate–phosphate) medium at 23°C under constant illumination of 5000 lux on a rotary shaker. Adult 2nd and 3rd leaves from *Z. mays* L. cultivar B73 was kindly provided by Dr. Lauri Smith (UC San Diego). Plants were grown in 4-inch pots in a greenhouse (temp: 23°C-29°C) without supplemental lighting or humidification (humidity in the 15 hours following inoculation ranged between 70 and 90%) year round in La Jolla, CA. RNA from *Z. mays* L. cultivar B73 7d old shoot, root and leaves were grown in the Schmitz laboratory (University of Georgia) as described in (Ricci et al. 2019).

### Data overview

A table of all generated and analyzed data can be found in Table S3

### csRNA-Seq Library Preparation

csRNA-seq was performed as described in (Duttke et al. 2019). Small RNAs of ∼20-60 nt were size selected from 0.4-3 µg of total RNA by denaturing gel electrophoresis. A 10% input sample was taken aside, and the remainder enriched for 5’-capped RNAs. Monophosphorylated RNAs were selectively degraded by 1 hour incubation with Terminator 5’-Phosphate-Dependent Exonuclease (Lucigen). Subsequently, RNAs were 5’dephosporylated through 90 minutes incubation in total with thermostable QuickCIP (NEB) in which the samples were briefly heated to 75°C and quickly chilled on ice at the 60 minutes mark. Input (sRNA) and csRNA-seq libraries were prepared as described in (Hetzel et al. 2016) using RppH (NEB) and the NEBNext Small RNA Library Prep kit, amplified for 11-14 cycles.

### Total RNA-Seq Library Preparation

Strand-specific, paired-end libraries were prepared from total RNA by ribosomal depletion using the Ribo-Zero Gold plant rRNA removal kit (Illumina, San Diego, CA). Samples were processed following the manufacturer’s instructions.

### Sequencing Information

Capped small RNA-seq libraries were sequenced on an Illumina NextSeq 500 instrument in the Benner lab or, like the total RNA-seq libraries, using a NovaSeq S6 at the IGM genomics core at UC San Diego. Information on read counts and alignment statistics can be found in Supplementary Tables S4.

## Data analysis

List of used genomes and annotations:

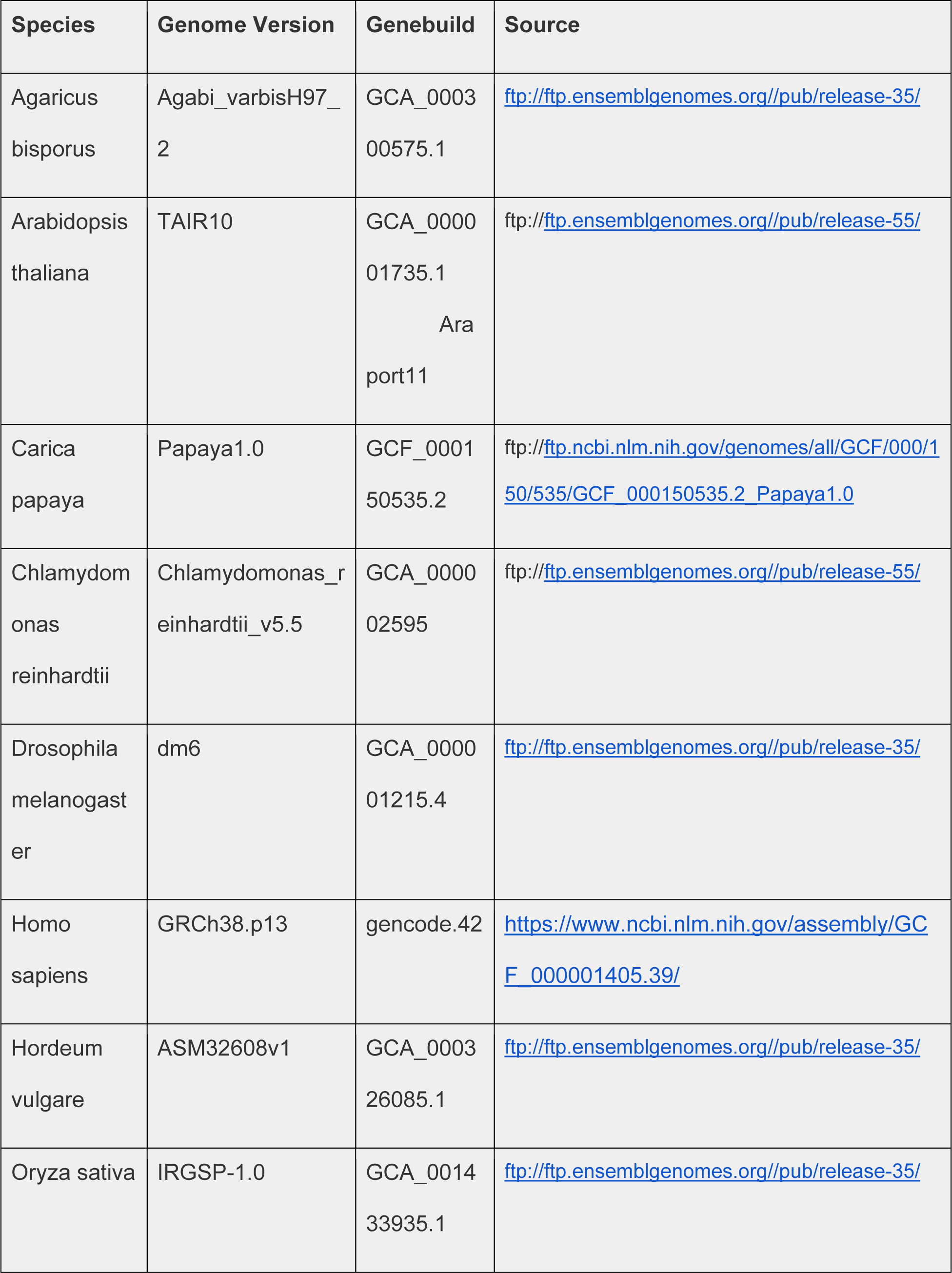

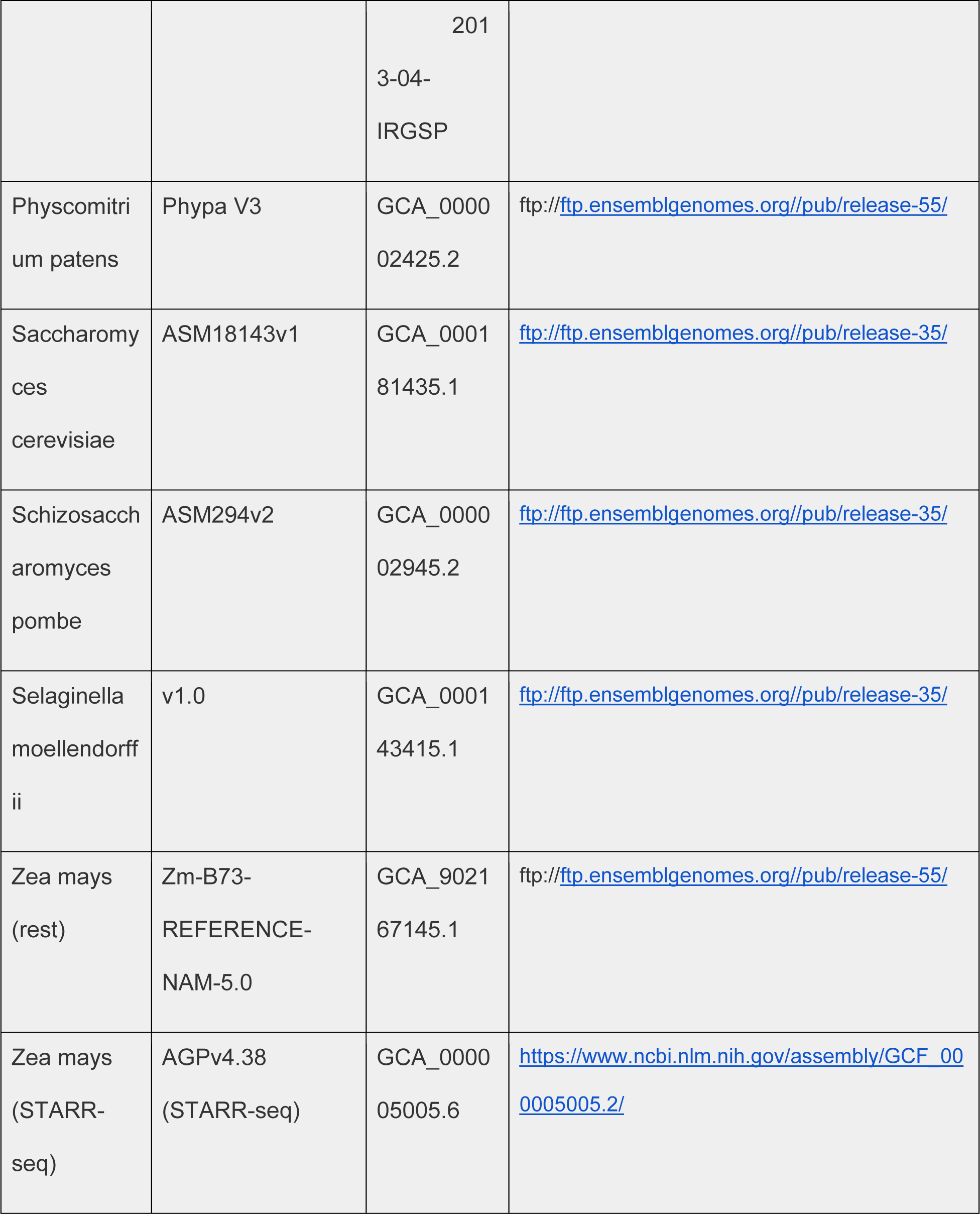

### csRNA-seq Data Analysis

Transcription start regions (TSRs), transcription start sites (TSSs) and their activity levels were determined by csRNA-seq and analyzed using HOMER v4.12 (Duttke et al. 2019). Additional information, including analysis tutorials are available at https://homer.ucsd.edu/homer/ngs/csRNAseq/index.html. TSR files for each experiment were added to the GEO data.

csRNA-seq (small capped RNAs, ∼20-60 nt) and total small RNA-seq (input) sequencing reads were trimmed of their adapter sequences using HOMER (“batchParallel.pl “homerTools trim -3 AGATCGGAAGAGCACACGTCT -mis 2 -minMatchLength 4 -min 20” none -f {csRNA_fastq_path}/*fastq.gz”) and aligned to the appropriate genome using *Hisat2* (Kim et al. 2019) (“hisat2 -p 30 --rna-strandness RF --dta -x {hisat2_genome_index} -U {path_rimmed_csRNA or sRNA} -S {output_sam} 2> {mapping_stats}”). Hisat2 indices were generated for each genome using “hisat2-build -p 40 genome.dna.toplevel.fa {Hisat2_indexfolder} except barely which required addition of “ --large-index”. Homer genomes were generated using “loadGenome.pl -name {Homer_genome_name} -fasta {species.dna.toplevel.fa} -gtf {species.gtf}”. Only reads with a single, unique alignment (MAPQ >=10) were considered in the downstream analysis. The same analysis strategy was also utilized to reanalyze previously published TSS profiling data to ensure the data was processed in a uniform and consistent manner, with exception of the adapter sequences, which were trimmed according to each published protocol. Tag Directories were generated as described in the csRNA-seq tutorial. We automated the process for all species by first generating an infofile.txt and then generating them in a batch

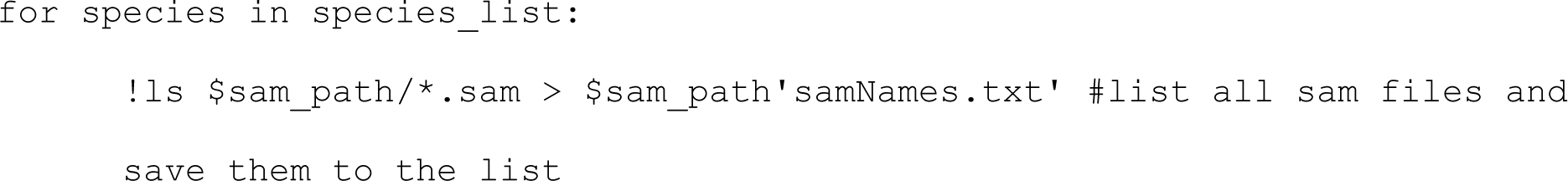
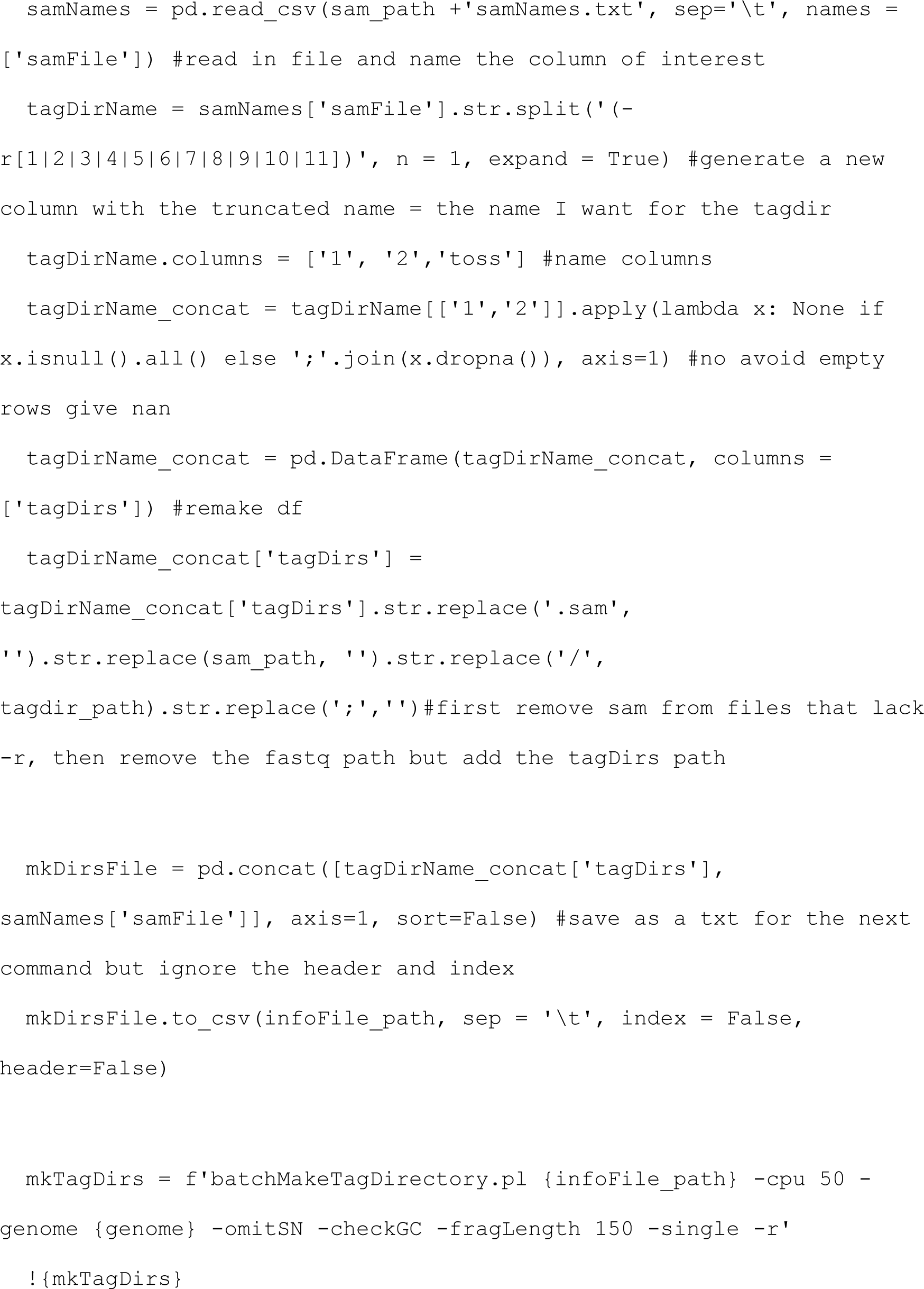

TSS and TSRs were analyzed in this study. TSR, which comprise one or several closely spaced individual TSS on the same strand from the same regulatory element (i.e. ‘peaks’ in csRNA-seq), were called using findcsRNATSS.pl (Duttke et al. 2019) (“findcsRNATSS.pl {csRNA_tagdir} -o {output_dir} -i {sRNA_tagdir} -rna {totalRNA_tagdir} -gtf {gtf} -genome {genome} -ntagThreshold 10”). findcsRNATSS.pl uses short input RNA-seq, total RNA-seq (Ribo0), and annotated gene locations to find regions of highly active TSS and then eliminate loci with csRNA-seq signal arising from non-initiating, high abundance RNAs that nonetheless are captured and sequenced by the method (for more details, please see (Duttke et al. 2019)). Replicate experiments were first pooled to form meta-experiments for each condition prior to identifying TSRs. Annotation information, including gene assignments, promoter distal, stable transcript, and bidirectional annotations are provided by findcsRNATSS.pl. To identify differentially regulated TSRs, TSRs identified in each condition were first pooled (union) to identify a combined set of TSRs represented in the dataset using HOMER’s mergePeaks tool using the option “-strand”. The resulting combined TSRs were then quantified across all individual replicate samples by counting the 5’ ends of reads aligned at each TSR on the correct strand. The raw read count table was then analyzed using DESeq2 to calculate normalized rlog transformed activity levels and identify differentially regulated TSRs (Love et al. 2014).

TSS were called using getTSSfromReads.pl (“getTSSfromReads.pl -d {csRNA_tagdir} -dinput {sRNA_tagdir} -min 7 > {output_file}” (Duttke et al. 2019)). To ensure high quality TSSs, at least 7 per 10^7 aligned reads were required and TSS were required to be within called TSR regions (subsequently filtered using mergePeaks “mergePeaks {TSS.txt} {stableTSRs.txt} -strand - cobound 1 -prefix {stable_tss}” or “mergePeaks {TSS.txt} {unstableTSRs.txt} -strand -cobound 1 -prefix {unstable_tss}”). Furthermore, TSSs that had higher normalized read density in the small RNA input sequencing than csRNA-seq were discarded as a likely false positive TSS location.

These sites often include miRNAs and other high abundance RNA species which are not entirely depleted in the csRNA-seq cap enrichment protocol. In most cases, TSRs were analyzed (i.e. to determine motifs or describe the overall transcription activity of regulatory elements) but when indicated single nucleotide TSS positions were independently analyzed (i.e. to determine motif spacing to the TSS).

Annotation of TSS/TSR locations to the nearest gene was performed using HOMER’s annotatePeaks.pl program using GENCODE as the reference annotation (Heinz et al. 2010).

For additional information about csRNA-seq analysis and tips for analyzing TSS data, please visit the HOMER website: https://homer.ucsd.edu/homer/ngs/csRNAseq/index.html. Strand specific and other IGV and genome browser files were generated using “makeUCSCfile {tag_directory_name} -strand + -fragLength 1 -o {tag_directory_name}.bedGraph” where the tag_directory could be csRNA-seq or 5’GRO-seq data from any species or tissue.

### 5’GRO-seq and GRO-seq analysis

Published and generated 5’GRO-seq and GRO-seq data were analyzed as described for csRNA-seq and sRNA-seq above. 5’GRO-seq peaks were called using HOMER’s “findPeaks {5GRO_tagdirectory} -i {GRO_tagdirectory} -style tss -F 3 -P 1 -L 2 -LP 1 -size 150 -minDist 200 -ntagThreshold 10 > 5GRO_TSRs.txt”. Detailed explanation of each parameter can be found at http://homer.ucsd.edu/homer/ngs/tss/index.html.

### RNA-seq analysis

Paired end total ribosomal RNA-depeleted RNA-seq libraries were trimmed using skewer (“time -p skewer -m mp {read1} {read2} -t 40 -o {trimmed_fastq_output}”) (Jiang et al. 2014) and aligned using *Hisat2* (Kim et al. 2019) to ensure all data were processed as similar as possible (“hisat2 -p 30 --rna-strandness RF --dta -x {hisat2_index} -1 {trimmed_RNAseq_R1} -2 {trimmed_RNAseq_R2} -S {output_sam} 2> {mapping_file}”). In this manuscript, total RNA-seq was exclusively used to determine RNA-stability as described in the csRNA-seq analysis.

### ChIP-seq analysis

Tag directories were generated for paired end sequenced ChIP libraries as described for total RNA-seq. Peaks were called using HOMER’s **“**findPeaks {ChIP_tagdir} -i {ChIP_inout_tagdir} - region -size 150 -minDist 370 > ChIP_peaks.txt” Quantification of histone modifications associated with each TSS was performed from +1 to +600 to capture the signal located just downstream from the TSS. When reporting log2 ratios between read counts a pseudocount of “1 read” was added to both the numerator and denominator to avoid divide by zero errors and buffer low intensity signal.

### ATAC-seq analysis

ATAC-seq data were analyzed as described for csRNA-seq but trimmed using “CTGTCTCTTATACACATCT”.

### DNA Motif Analysis

*De novo* motif discovery and motif occurrence scanning were performed using HOMER (Heinz et al. 2010)using TSRs as input. Binding sites were searched from -150 to +50 bp relative to the primary TSS from each TSR, using GC-matched random genomic regions as background. Motif occurrences, including histograms showing the density of binding sites relative to the TSS (reported as motifs per bp per TSS), and average nucleotide frequency were calculated using HOMER’s annotatePeaks.pl tool. Known motifs were analyzed using findMotifsGenome.pl with the option -mknown to specifically analyze a motif or sets of motifs including our plant DNA sequence motif library. For core promoter elements including the TATA-box or the Initiator, the area where the motif is searched were constrained to their respective preferences relative to TSSs. For example, TSSs with a match to BBCA+1BW (where the A+1 defines the initiating nucleotide) were considered Inr-containing TSS (Vo ngoc et al. 2017). The “-norevopp” function was used to make the search strand specific and “-size -6,6” was used for Inr motifs, “ -size - 35,20” for TATA-box and TATA-box-like motifs. Data were summarized using custom python code, transformed using pandas melt function (“pd.melt(CPE_frame, id_vars=[’species’], value_vars=[’TATA’, ’TATA_1mm’, ’hINR’, ’dINR’])”. https://zenodo.org/record/7794821#.ZEAz_XbMJ3g) and plotted using seaborn (“sns.boxplot”) [10.21105/joss.03021]. Additional information can be found on the HOMER website: http://homer.ucsd.edu/homer/motif/.

To identify de novo motifs enriched in the TSRs of all plant species and tissues, sequences of each TSR were combined into one fasta file and findMotifs.pl used. The motif search was constrained to sequences of 8 bp length to focus on the motif core and sequences 2000 bp downstream of the TSR selected as background (“findMotifs.pl all_plantTSRs.fa fasta outputdir/-fasta all_plantTRSbackground.fa -len 8 -S 200 -noconvert -p 60”). Subsequently, all known plant motifs in the updated HOMERplants motif library (Hetzel et al. 2016) were added and redundant motifs removed using “compareMotifs.pl {input.motif} {output_folder} -cpu 40 - reduceThresh 0.6” and subsequently non-similar motifs concatenated “cat output_folder/homerResults/motif[!V].motif output_folder/homerResults/motif?[!V].motif output_folder/homerResults/motif??[!V].motif > NonRedudnant06.motif”.

For the Motif enrichment plot in each species (Fig.2B), the top 5 motifs from each species or tissue were combined (“combineGO.pl -top 5 -f knownResults.txt -d {homerresults_each_species}/* > all_species_5_motifs.txt”) and plotted using sns.clustermap.

### Motif correlation of stable and unstable TSRs

Motifs were defined using HOMER and our 151 motif library using stable or unstable TSRs as foreground and the other as background (“findMotifsGenome.pl {stable_TSS_file} {species_fa} {species_tss}_stable/ -bg {UNstable_TSS_file} -mask -p 40 -size -150,50 -mset all -S 15 -len 10 findMotifsGenome.pl {UNstable_TSS_file} {species_fa} {species_tss}_UNstable/ -bg {stable_TSS_file} -mask -p 40 -size -150,50 -mset all -S 15 -len 10”). Frames were concatenated and the correlation calculated using pandas .corr function (https://zenodo.org/record/7794821#.ZD1rA3bMKUk).

### Transcript stability switch analysis

Transcript stability was determined as unstable if <2/10^7 total RNA-seq reads were within - 100,+500 of the main TSS of the TSR. In *Arabidopsis* we compared cells and adult leaves to identify transcripts that had differential stability among the conditions, in maize we used adult leaves, 7d-old seedling leaves, 7d-old seedling roots and 7-d old seedling shoots. For the plots (Fig. 5d; “sns.pointplot”) we limited our analysis in maize to 7d-old shoot vs. root.

### Mapping stats calculation

All 2> outputs from hisat2 were copied into a mappingstats folder and summarized using the following custom code.

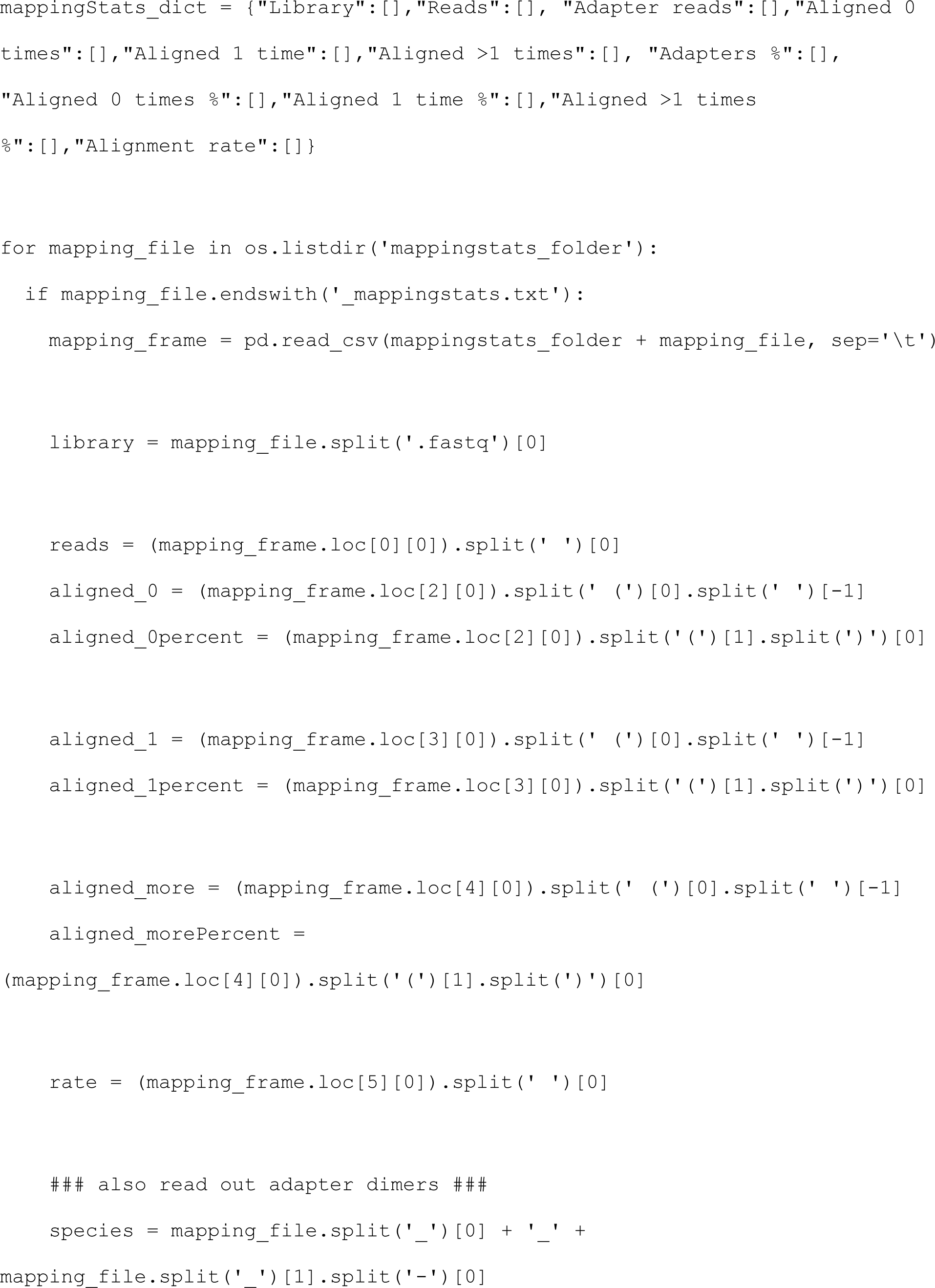
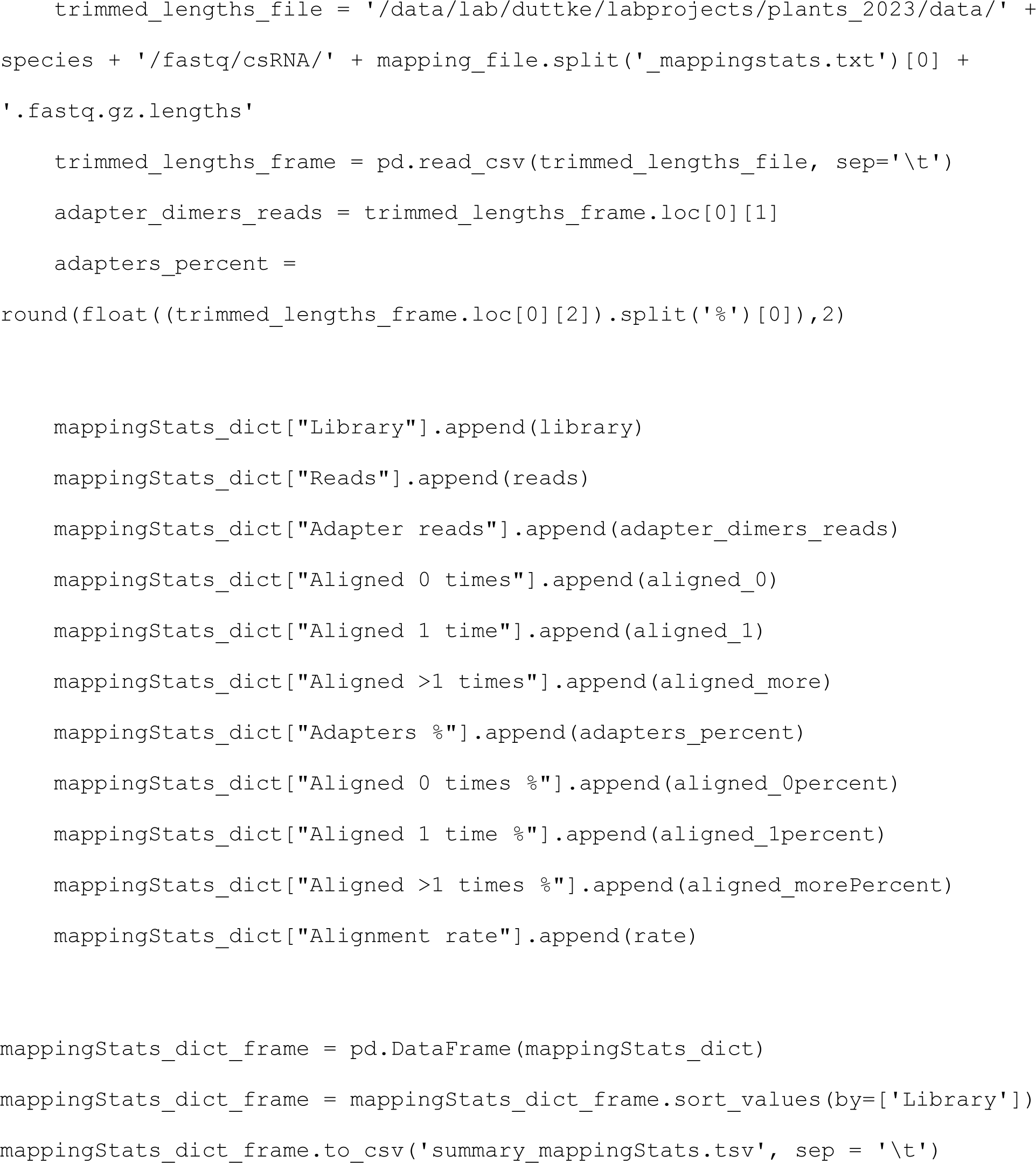

### Histograms and annotation of TSS to captured reads

Histograms showing csRNA-seq or other data relative to known TSS were generated using “annotatePeaks.pl {known TSS} {species_homer_genome (e.g. TAIR10)} -strand + -fragLength 1 -size 100 -d {species_tagdirectory (e.g. P.patens_csRNAseq)} -raw > output.tsv. “Known TSS” were extracted from .gtf files using “parseGTF.pl {species_gtf_file} tss > {species}_genes.tss”. Histograms showing called TSS by csRNA-seq or 5’GRO-seq relative to one another or “Known TSS” were generated using “annotatePeaks.pl {reference or “Known TSS} {species_homer_genome (e.g. TAIR10)} -p {2nd TSS file, i.e. csRNA-seq TSS} -size 2000 -hist 1 -strand + > output.tsv”.

### Gene Ontology Analysis

Gene Ontology analysis was performed using METASCAPE (Zhou et al. 2019) for transcripts annotated within 500 bp downstream of the main TSS of TSRs.

### STARR-seq analysis

csRNA-seq data were generated from analogous tissue as used for STARR-seq (GSE120304_STARR_B73_enhancer_activity_ratio.txt.gz) by Ricci et al (2019) as described above. For compatibility reasons, this analysis thus used the *Z. mays* AGPv4 reference genome and *Z. mays* AGPv4.38 genome annotation) instead of maize 5.5. STARR-seq library fragments of 1-50 bp were removed from the analysis as these short fragments disproportionally showed no enhancer activity while longer fragments of the same locus did. csRNA-seq TSRs were defined as described above and merged with the STARR-seq peaks (“mergePeaks”) to identify overlaps. As sometimes several STARR-seq peaks fell within on TSR, we next corrected the STARR-seq values by linking each mergedPeak ID with the sum of STARR-seq peaks that fell within the peak. Next we normalized this value by the length of the peak to obtain a STARR-seq value per bp for each merged peak and added the csRNA-seq values and TSR stability.

### Pausing index

The pausing index was calculated as described (Chen et al. 2015) using reads near TSRs (- 100bp to +300bp) divided by those found downstream in the region of +301 to +2kb, relative to the major TSS of the TSR.

## Data availability

All raw and processed data generated for this study can be accessed at NCBI Gene Expression Omnibus (GEO; https://www.ncbi.nlm.nih.gov/eo/) accession number GSE233927. Reviewer access token: mtsxkwewnlsvxep

## Code availability

Code used to analyze data in this manuscript has been described in the methods, or is available from the following repositories:

HOMER (http://homer.ucsd.edu/)

MEIRLOP (https://github.com/npdeloss/meirlop)

## Acknowledgments

We thank Dr. Ashley M Brooks and Drs. Laurie Smith, Will Ansari, Pete Hedley, Stephen Mayfield for generous donation of plant tissues and Dr. Tamar Juven-Gershon for *Drosophila* S2 cell RNA. We thank Emma L. Ledbetter and Life Science Editors for manuscript editing, Christopher W. Benner for help with data generation, James T. Kadonaga, Mackenzie K. Meyer, and Jacob W. Bonner for useful discussion. B.R.M is a STARS undergraduate fellow. This work was supported by NIH grants R00GM135515 to S.H.D. R.J.S. was supported by the National Science Foundation (IOS-1856627). S.E.J is an Investigator of the Howard Hughes Medical Institute. This publication includes data generated at the UC San Diego IGM Genomics Center utilizing an Illumina NovaSeq 6000 that was purchased with funding from a National Institutes of Health SIG grant (#S10 OD026929)

## Author Contributions

R.J.S, S.E.J., and S.H.D. oversaw the overall design and execution of the project. The experiments were performed by B.R.M., I.M.B., M.I.S. and S.H.D. The computational analyses were performed by B.R.M., I.M.B., C.P., and S.H.D. C.P and S.H.D. were primarily responsible for writing the manuscript. All authors revised and approved the final manuscript.

## Corresponding author

Correspondence to Sascha H. Duttke

## Ethics declarations

The authors declare no competing interests.

